# Polymer-based flexible multi-shank probes for simultaneous intracortical microstimulation and two-photon calcium imaging

**DOI:** 10.64898/2026.02.09.704741

**Authors:** Eszter Nguyen, Balázs Barkóczi, Csaba Horváth, Linda Judák, Balázs Rózsa, Frederik Ceyssens, István Ulbert, Lucia Wittner, Maarten Schelles, Richárd Fiáth

## Abstract

Intracortical microstimulation is an essential tool for basic neuroscience and sensory restoration, yet the spatiotemporal effects of advanced multielectrode stimulation strategies on cortical networks remain poorly understood. This study presents a versatile experimental platform featuring polymer-based, flexible multi-shank electrode arrays integrated with a custom high-density neurostimulator. The polyimide penetrating probes contain densely spaced iridium oxide microelectrodes on thin, flexible shanks with sharp tips designed to minimize mechanical mismatch, tissue damage, and brain dimpling. The system was validated in transgenic GCaMP6 mice through simultaneous two-photon calcium imaging and electrical stimulation in layer 2/3 of the visual cortex. Monopolar stimulation reliably evoked robust, parameter-dependent neuronal activation, with neuronal recruitment increasing as a function of current intensity and pulse duration. Bipolar stimulation produced activation patterns distinct from monopolar stimulation, with neuronal responses that were more spatially confined when bipolar electrodes were positioned in close proximity. Furthermore, static current steering—implemented by varying the current ratio between the selected electrode pair—successfully shifted the centroid of activated neuronal populations, enabling fine spatial control over the site of activation. The developed platform provides a flexible framework for investigating local neuronal responses to complex electrical stimulation paradigms, facilitating the development of neural prostheses with higher spatial precision.

## 1. Introduction

Electrical stimulation is one of the most widely used neuromodulation techniques in both basic research and clinical applications. In animal models, it is commonly employed to study its effects on behaviour and to investigate stimulation-induced cellular and network-level changes in neuronal activity (Bari et al., 2013; Cheney et al., 2013; Clark et al., 2011; Dadarlat et al., 2024; Eles & Kozai, 2020; Histed et al., 2009; Histed et al., 2013; Hughes et al., 2025; McIntyre & Grill, 2000; Meikle et al., 2022; Michelson et al., 2019; Ozen et al., 2010; Stieger et al., 2020, 2022; Tehovnik et al., 2006). In humans, electrical stimulation is applied to diagnose or treat a range of central and peripheral nervous system disorders. Examples include deep brain stimulation for alleviating the symptoms of Parkinson’s disease (Krauss et al., 2021; Lozano et al., 2019; Perlmutter & Mink, 2006) and sensory implants (e.g., retinal, cochlear or somatosensory prostheses) for restoring sensory function (Beauchamp et al., 2020; Bosking et al., 2017; Carlyon & Goehring, 2021; Cicmil & Krug, 2015; Lewis et al., 2015; Shepherd et al., 2013; Shim et al., 2020; Tehovnik & Slocum, 2013a; Tyler, 2015; Zeng et al., 2008). Intracortical microstimulation (ICMS) is a specific form of electrical stimulation in which currents in the microampere range are delivered into cortical tissue with penetrating neural interfaces to modulate neuronal firing or to elicit sensory percepts (Chen et al., 2020; Eles & Kozai, 2020; Flesher et al., 2016; Lycke et al., 2023; Stieger et al., 2022).

Most current research protocols and clinical applications rely on monopolar stimulation, where the current flows between a single active microelectrode and a distant ground (Cunningham et al., 2025; Histed et al., 2009; Michelson et al., 2019; Zhu et al., 2012). Additionally, these approaches typically use only a small number of electrodes for stimulation. However, monopolar stimulation often produces widespread neuronal activation and offers limited control over the electric field compared with multipolar or more advanced stimulation approaches and therefore lacks the spatial selectivity required for many applications (Moleirinho et al., 2021; Spencer et al., 2016; Zhu et al., 2012). For instance, achieving high-resolution vision with retinal or cortical implants would require stimulating small, spatially confined neuronal populations corresponding to functional units of vision, such as cortical columns (Christie et al., 2016; Jepson et al., 2014; Tong et al., 2020). For monopolar stimulation, this would demand an impractically large number of densely packed and miniature electrodes, which is currently not feasible in the retina and remains highly challenging in the human visual cortex. Consequently, alternative stimulation strategies are needed to achieve high-spatial-resolution activation using fewer implanted electrodes (Dumm et al., 2014; Jepson et al., 2014; Lyu et al., 2020; Opie et al., 2013; Tong et al., 2020).

Multielectrode stimulation strategies—including bipolar stimulation, in which current flows between two closely spaced electrodes of opposite polarity; current steering, where currents delivered through multiple electrodes are independently adjusted to shape the electric field; and current focusing, in which coordinated stimulation across electrodes confines current spread—enable controlled shaping of the resulting electric field and promote more selective activation of targeted neuronal populations (Berenstein et al., 2008; Bonham & Litvak, 2008; Dumm et al., 2014; Meikle et al., 2022; Opie et al., 2013; Spencer et al., 2018; Uguz & Shepard, 2022; Zhu et al., 2012). Such approaches are already used in clinical applications, including deep brain stimulation and cochlear implants, where steering and focusing the electric field between electrodes enhances spatial selectivity (Berenstein et al., 2008; Bonham & Litvak, 2008; Chaturvedi et al., 2012; Slopsema et al., 2018; Zhu et al., 2012).

However, despite growing interest in these paradigms, their effects on ongoing cortical activity remain poorly understood. Several factors contribute to this knowledge gap. First, electrophysiological techniques are the most widely used methods in this field; however, electrical stimulation introduces substantial artifacts into electrophysiological recordings, complicating the analysis and interpretation of spatiotemporal response patterns. Second, extracellular recordings typically capture spiking activity only from neurons located close (< 150 µm) to the recording electrode, and accurately determining neuronal positions relative to the stimulation electrode is inherently challenging. Moreover, most microelectrodes used for stimulation have relatively large surface areas and therefore predominantly record multi-unit activity due to signal averaging, limiting access to single-neuron responses. Third, integrated hardware and software platforms for multielectrode intracortical microstimulation that allow flexible adjustment of stimulation parameters and electrode selection to evaluate the local effects of different electrical stimulation strategies are not widely available. Fourth, most penetrating stimulation electrodes are fabricated from rigid materials (e.g., tungsten or silicon), which exhibit a substantial mechanical mismatch with soft brain tissue. This mismatch can lead to significant neuronal loss, inflammation and glial scar formation over time in the vicinity of the stimulation site, thereby distorting physiological neural responses to electrical stimulation in long-term experiments (McConnell et al., 2009; Polikov et al., 2005; Potter et al., 2012).

Many of these challenges can be addressed by combining electrical stimulation with alternative recording methods and by optimizing electrode materials. For instance, optical imaging techniques, such as two-photon calcium imaging, allow simultaneous observation of hundreds of neurons during stimulation in larger tissue volumes, with precise spatial localization and without stimulation-induced recording artifacts (Grienberger et al., 2022; Tehovnik & Slocum, 2013b). Additionally, neural probes fabricated from softer, flexible materials (e.g., polyimide, SU-8, or Parylene C) may mitigate neuronal damage and the foreign body response, enabling more reliable results in chronic studies (He et al., 2020; Lycke et al., 2023; Orlemann et al., 2024; Zhao et al., 2023; Zhou et al., 2017).

In this study, we developed a versatile neurostimulator system together with polymer-based, flexible multi-shank probes featuring densely spaced microelectrodes, providing a flexible experimental framework to investigate local neuronal responses to diverse electrical stimulation paradigms. Micro-electromechanical systems (MEMS) technology enables the fabrication of probes with arbitrary microelectrode configurations, while the flexibility and small cross-sectional dimensions of the polyimide shanks reduce both acute and chronic tissue damage, indirectly improving the quality of optical imaging. We validated the functionality of the system by performing monopolar stimulation across a range of stimulation parameters in anesthetized and awake, head-fixed transgenic GCaMP6 mice, with microelectrodes implanted in layer 2/3 of the visual cortex. Stimulation-evoked neuronal responses were obtained using two-photon microscopy over a field of view of 550 µm × 550 µm at a frame rate of approximately 31 Hz. The electrophysiological performance of the electrode arrays was also characterized. Finally, we applied bipolar stimulation and current steering in anesthetized mice to assess the capabilities of the system for multielectrode current delivery.

## 2. Results and Discussion

### 2.1. Flexible multi-shank probe design

We designed and fabricated polyimide-based penetrating probes with multiple, flexible shanks containing closely packed microelectrodes, along with a dedicated stimulator device, with the aim of providing a versatile experimental platform for studying how distinct electrical stimulation paradigms modulate local cortical activity (Figure 1a,b). Platinum interconnects linking the bonding pads to iridium oxide microelectrodes are embedded between two 10-µm-thick polyimide layers (Figure 1c). Each 1 mm-long flexible shank contains six rectangular microelectrodes (size: 50 µm × 30 µm; electrode pitch: 75 µm) and one triangular electrode forming a sharp tip at the distal end of the shank. Two probe geometries were implemented, with intershank pitches of 325 µm and 650 µm for the three- and two-shank designs, respectively, specifically designed to enable systematic investigation of local neuronal responses to electrical stimulation delivered between microelectrodes separated by short (<1 mm) distances (Figure 1a,b). To facilitate probe handling during implantation, a small and rigid SU-8 support structure is integrated at the probe base above the shanks (Figure 1a,b). In addition, the electrode array features a long, meandering cable section designed to reduce mechanical strain between the implanted shanks and the external connector (Figure 1a,b).

**Figure 1.**
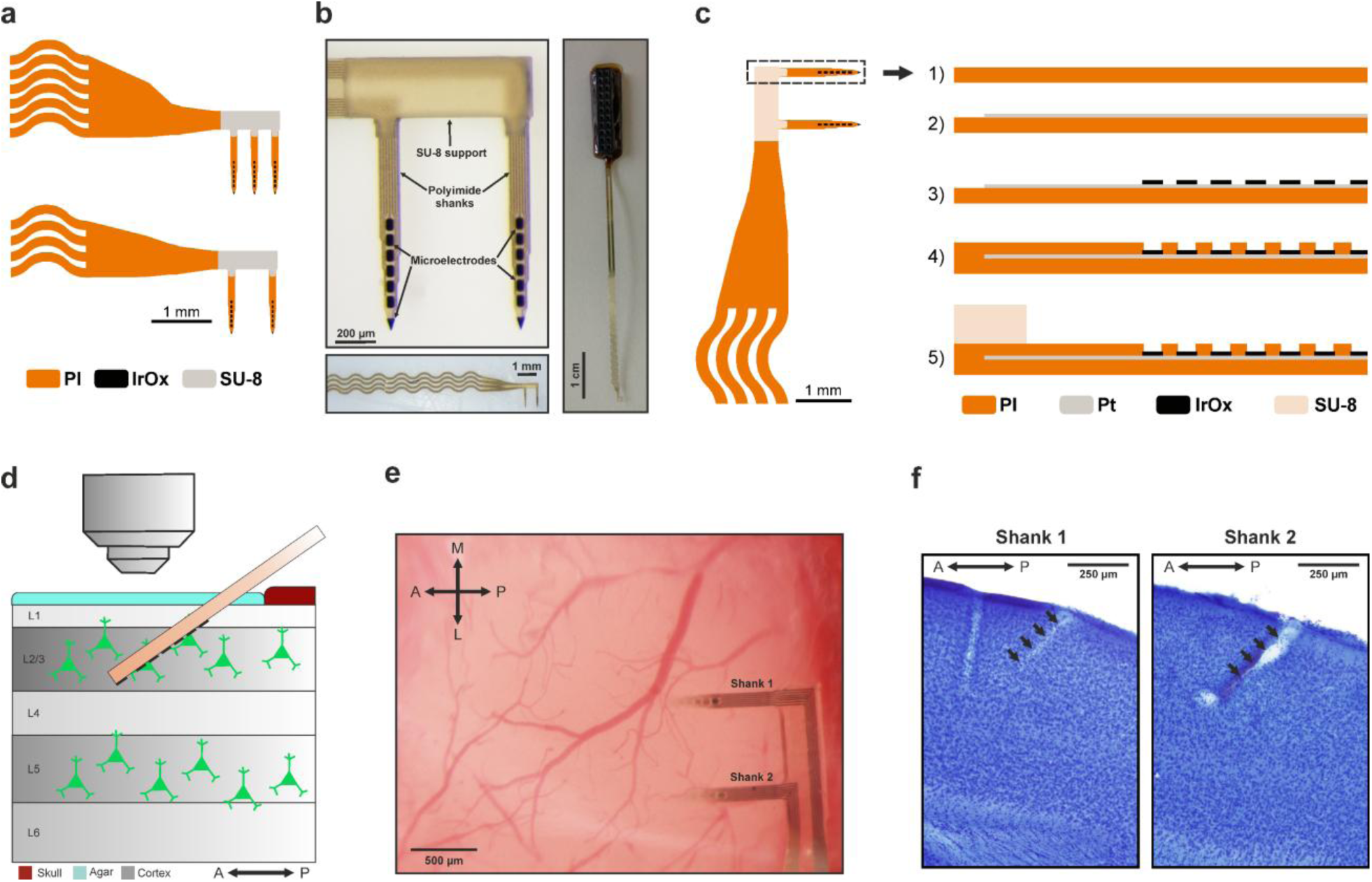
Probe design and experimental setup. a) Schematic of the multi-shank probes. b) Stereomicroscopic images of the two-shank (left) and three-shank (right) probe variants at different magnifications. The probe comprises a long, meandering cable section (bottom left). c) Main steps of the probe fabrication process. IrOx, iridium oxide; PI, polyimide; Pt, platinum. d) Schematic side view of the experimental setup. Probes were inserted into the primary visual cortex at a 55° angle from vertical to a depth of ∼300 µm from the cortical surface, targeting the superficial cortical layers. Calcium imaging was performed in the vicinity of the probe shanks using a two-photon laser scanning microscope. e) Stereomicroscopic top view image of a two-shank probe implanted into the visual cortex. f) Nissl-stained sagittal brain sections showing the two insertion tracks (black arrows) of a two-shank probe in the visual cortex. The tracks correspond to the implantation shown in panel e. A, anterior; M, medial; L, lateral; P, posterior.

The flexible probes were designed for simultaneous electrical stimulation and *in vivo* calcium imaging in mice. In the *in vivo* experiments presented here, the electrode arrays were implanted into the superficial layers of the primary visual cortex (V1) of GCaMP6-expressing transgenic mice and cortical activity was monitored within a 550 µm × 550 µm field of view (FOV) using two-photon laser scanning microscopy (Figure 1d). To minimize tissue damage and microbleedings during insertion, probes were advanced at a slow rate (∼2 µm/s) using a remote-controlled micro-tweezer tool and a motorized stereotaxic setup (Fiath et al., 2019). This implantation approach typically preserved vascular integrity at the insertion site (Figure 1e) and resulted in thin, often barely detectable insertion tracks, as confirmed by post-hoc histology (Figure 1f). Minimizing brain dimpling is also essential for achieving high-quality calcium imaging in superficial cortical layers. The sharp-tipped shank design contributed to low dimpling, which could be further reduced by making small incisions in the dura mater at the shank entry points.

### 2.2. *In vitro* characterization of flexible probes

Although the electrode array was primarily designed for stimulation, it is inherently capable of recording brain electrical activity, including local field potentials (LFP; < 500 Hz) and single-/multi-unit activity (SUA/MUA; > 500 Hz). The electrophysiological capability of the probe is particularly valuable for multimodal approaches, such as combining optical imaging with electrophysiology, or for simultaneous electrical stimulation and neural recording.

To evaluate the stimulation and electrophysiological recording performance of the fabricated devices, we first performed *in vitro* electrical impedance spectroscopy on 77 electrodes across four probes (Figure 2a). The impedance magnitude of the iridium oxide microelectrodes (1500 µm^2^ area) was found on average 17.32 kΩ ± 4.96 kΩ measured at 1 kHz, which is suitable for low-noise spike recordings (Figure 2a). Only two microelectrodes were found to be nonfunctional, exhibiting high (> 1 MΩ) impedance values.

**Figure 2.**
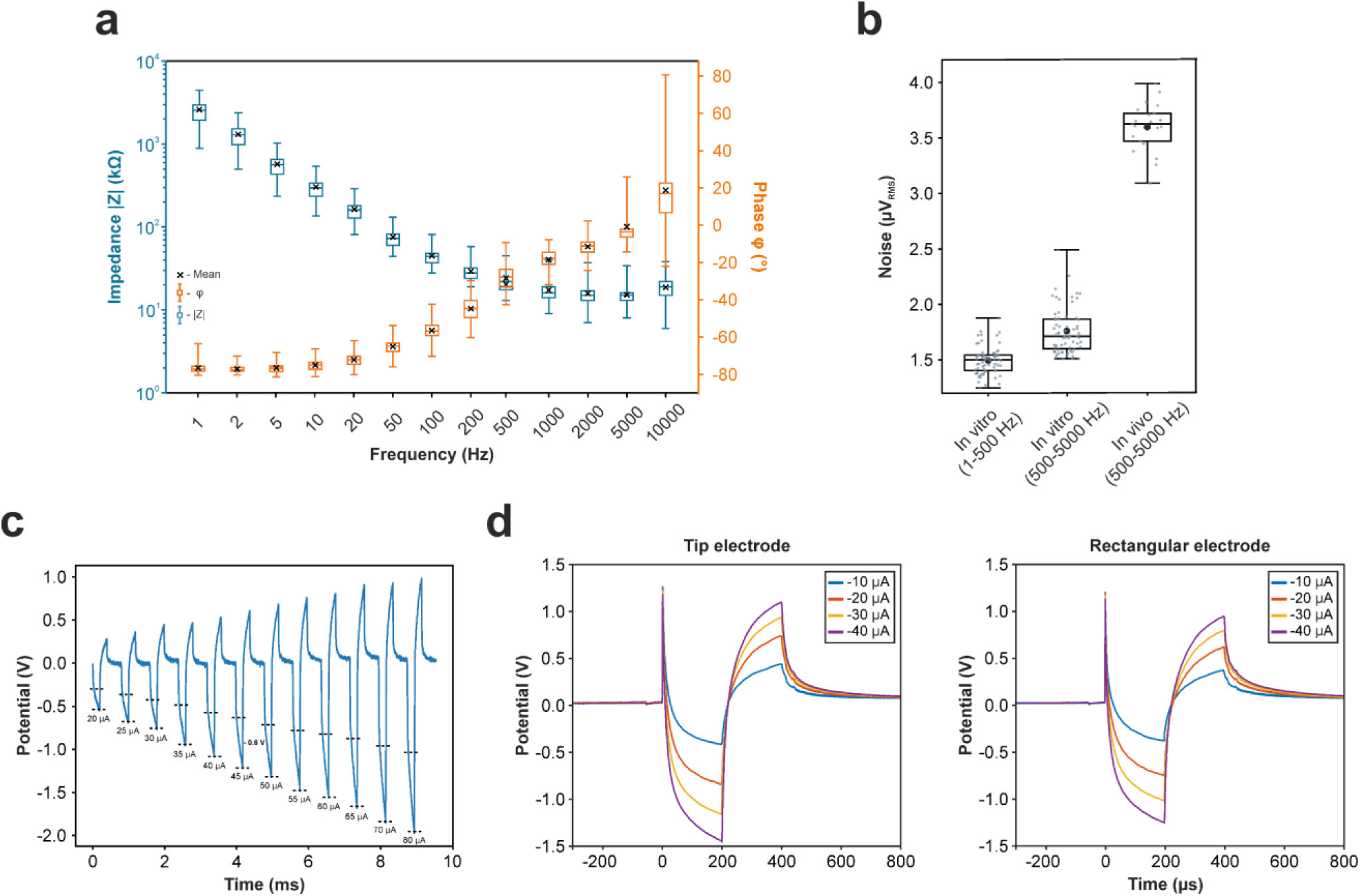
In vitro characterization of flexible probes. a) Box-whisker plots showing the impedance magnitude (blue) and phase (orange) of 75 iridium-oxide microelectrodes (n = 4 probes; data from two nonfunctional electrodes were excluded) measured at 13 frequencies. b) Box-whisker plots of in vitro (n = 67 microelectrodes) and in vivo (n = 21 electrodes) root mean square (RMS) noise levels. In vitro RMS noise was measured in both the local field potential (1 - 500 Hz) and unit activity (500 - 5000 Hz) bands. c) Representative example of in vitro measurements of the charge injection capacity of the electrodes. Pulses started at an amplitude of 20 µA, and increased with steps of 5 µA. The cathodic capacitive voltage drop of each pulse is indicated with dashed lines. d) Representative voltage waveforms of stimulation pulses acquired at four current intensities for a triangular tip electrode (left) and a rectangular microelectrode (right). Each trace shows the mean of 100 individual pulses.

Next, we quantified *in vitro* and *in vivo* noise levels. *In vitro* root-mean-square (RMS) noise was 1.50 µV ± 0.13 µV and 1.76 µV ± 0.20 µV in the LFP (1 – 500 Hz) and SUA/MUA (500 – 5000 Hz) bands, respectively (Figure 2b). *In vivo* RMS noise levels, measured in the visual cortex of an anesthetized wild-type mouse, were slightly higher, with an average of 3.62 µV ± 0.21 µV (Figure 2b).

To assess stimulation performance, we measured the *in vitro* charge injection capacity of the electrodes, which was 0.57 mC/cm² ± 0.05 mC/cm². This corresponded to a safe stimulation current of 40-45 µA, before the water window was crossed (Figure 2c). Representative examples of charge-balanced, symmetric, biphasic, cathodic-leading voltage waveforms (averaged over 100 pulses, each with a duration of 400 µs) generated at different current intensities on a rectangular and a tip electrode are shown in Figure 2d.

### 2.3. *In vivo* electrophysiological recording capabilities of the flexible probe

As mentioned above, the developed devices are well-suited for multimodal experiments, such as those combining simultaneous calcium imaging and electrophysiological recordings. To assess their recording capabilities, probes were implanted into V1 of anesthetized C57BL6 mice to a depth of approximately 700 µm. We successfully recorded spontaneous high-quality local field potentials and spiking activity, capturing characteristic electrophysiological features of ketamine/xylazine-induced cortical slow waves, including an oscillation with a peak frequency of ∼1 Hz and the alternation of active (up) and silent (down) states with high and low neuronal firing, respectively (Figure 3a; Fiath et al., 2016; Neske, 2016). Most recording channels exhibited single-unit activity, with action potential amplitudes ranging from 48 µV to 210 µV (Figure 3b-d). Spike waveforms from individual neurons were frequently detected simultaneously on multiple adjacent electrodes (Figure 3c).

**Figure 3.**
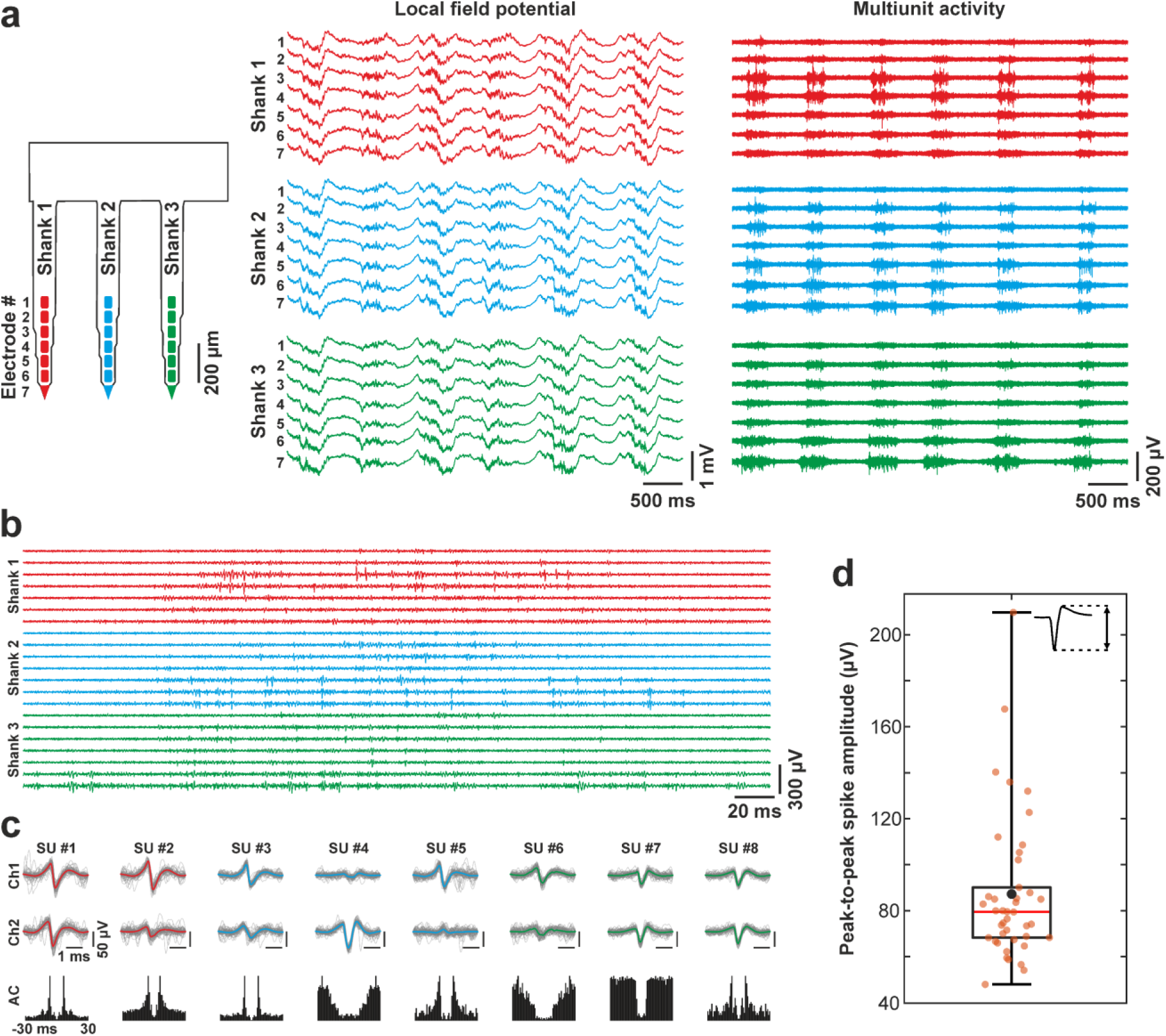
In vivo electrophysiological recording performance of the neural probe in anesthetized mice. a) Representative five-second-long traces of local field potential (1 - 500 Hz) and multi-unit activity (500 – 5000 Hz) recorded from the primary visual cortex of a mouse anesthetized with ketamine-xylazine. Channels corresponding to microelectrodes on different shanks of the probe are color-coded (see probe schematic on the left). b) Example recording snippet (∼300 ms; 500 – 5000 Hz band) showing single action potentials detected on most channels. c) Examples of isolated single units. Multichannel spike waveforms (top) and autocorrelograms (AC, bottom) are shown for eight units. For each unit, the mean spike waveform (color-coded according to the shank on which the unit was detected) is overlaid with 50 individual spike waveforms (gray). For each single unit, spike waveforms are shown for the two adjacent channels exhibiting the highest spike amplitudes (Ch1 and Ch2). d) Distribution of mean peak-to-peak spike amplitudes (see inset) for all single units detected in a single recording. Amplitudes were calculated from wideband mean spike waveforms.

Simultaneous electrophysiological recordings and calcium imaging provide complementary perspectives on brain activity (Higley & Cardin, 2022). While electrophysiology offers superior temporal resolution, two-photon calcium imaging enables precise spatial localization of active neurons (Siegle et al., 2021). Although direct imaging above microelectrodes induces strong high-frequency noise (photoelectric artifact) in the electrophysiological recordings (Orbán et al., 2019), electrodes located outside the imaging FOV can still capture high-quality cortical signals.

In an experiment conducted in a GCaMP6f mouse under light anesthesia, we performed calcium imaging around one of the probe shanks while simultaneously recording spontaneous cortical activity from the microelectrodes on the adjacent shank, located approximately 300 µm from the imaging FOV (Figure 4). Imaging was conducted at a depth of ∼400 µm below the cortical surface, corresponding roughly to layer 4, where several spontaneously active neurons were detected (Figure 4a).

**Figure 4.**
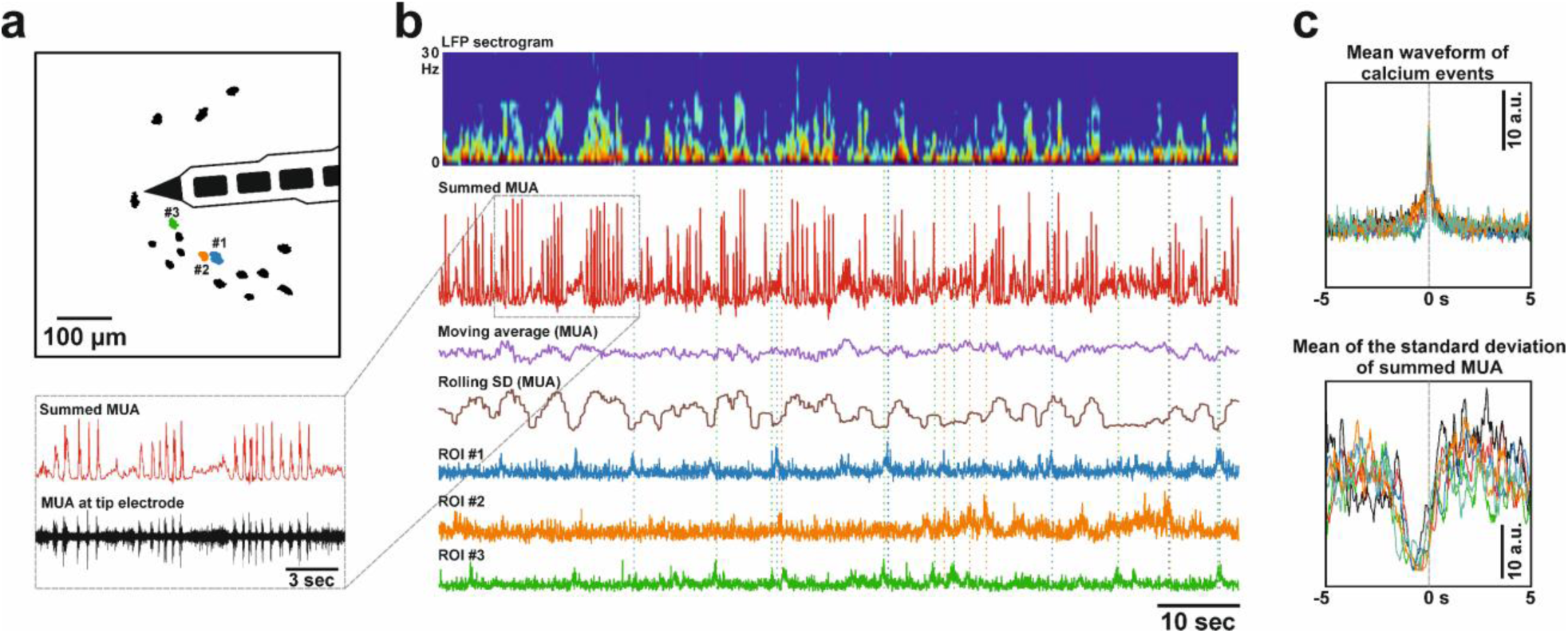
Simultaneous calcium imaging and electrophysiological recording in an anesthetized mouse. a) Schematic of the field of view (FOV) during calcium imaging around one probe shank, showing several regions of interest (ROIs). Neurons were located in layer 4 of the visual cortex. b) Short segment of spontaneous visual cortical activity obtained under ketamine/xylazine anesthesia. Electrophysiological signals (local field potential, LFP; multi-unit activity, MUA) were recorded from microelectrodes on the probe shank located outside the FOV shown in panel a, while calcium signals of the three ROIs were obtained within the FOV (see color-coded ROI masks). The LFP spectrogram and summed MUA (rectified and smoothed) reveal alternating short periods of synchronized and desynchronized neuronal activity (inset: synchronized periods in blue, desynchronized periods in green). This rhythmicity is evident in the standard deviation of the MUA (rolling SD) but not in the mean MUA (moving average). Neurons in the FOV preferentially fired during desynchronized periods, as highlighted by the colored vertical dashed lines indicating example calcium events for the three ROIs. c) Top: mean waveform of calcium events from six ROIs, aligned to the event peaks. Bottom: mean of the rolling SD of summed MUA, calculated around the calcium event times for the same ROIs. Signal variability is reduced before and during calcium events, suggesting that these neurons were most active during, or toward the end of, desynchronized periods, which are characterized by a lower standard deviation.

Analysis of both the LFP spectrogram and MUA summed across the seven electrodes outside the FOV revealed alternating short (few-second) periods of synchronized and desynchronized activity, with the synchronized periods—corresponding to slow waves and characterized by alternating up-and down-states—exhibiting greater amplitude variability (Figure 4b). Interestingly, neurons identified through calcium imaging tended to fire during, or toward the end of, desynchronized periods (Figure 4b, c). Although recorded MUA indicated that multiple neurons were active during the more rhythmic synchronized periods (most likely large layer 5 pyramidal cells, a key neuronal population involved in generating slow waves (Fiath et al., 2016), none of the imaged neurons displayed frequent calcium transients at those times (Figure 4b, c). These findings suggest that the two modalities - electrophysiology and calcium imaging - offer complementary insights into cortical network dynamics, providing a more complete picture of neuronal population activity than either modality alone.

### 2.4. *In vivo* calcium imaging in anesthetized mice during monopolar stimulation

To assess the *in vivo* stimulation performance of the developed probes and to validate overall system functionality, we implanted the electrode arrays into V1 of anesthetized GCaMP6 mice. Current-controlled monopolar stimulation was delivered between a distant ground electrode and a preselected rectangular or tip microelectrode positioned within the superficial layers (Figure 1d). The stimulation protocol consisted of a 10-s pre-stimulation (baseline) period, followed by a stimulation phase comprising several consecutive trials of pulse trains (Figure 5a). Each pulse train contained 40 or 100 symmetric, biphasic, cathodic-first pulses delivered at varying current intensities, pulse durations, and stimulation frequencies (see Methods for details). Individual stimulation trials were separated by 3-5 s intervals, and the whole stimulation protocol (during which all stimulation parameters were held constant) concluded with a post-stimulation period lasting approximately 10 s.

**Figure 5.**
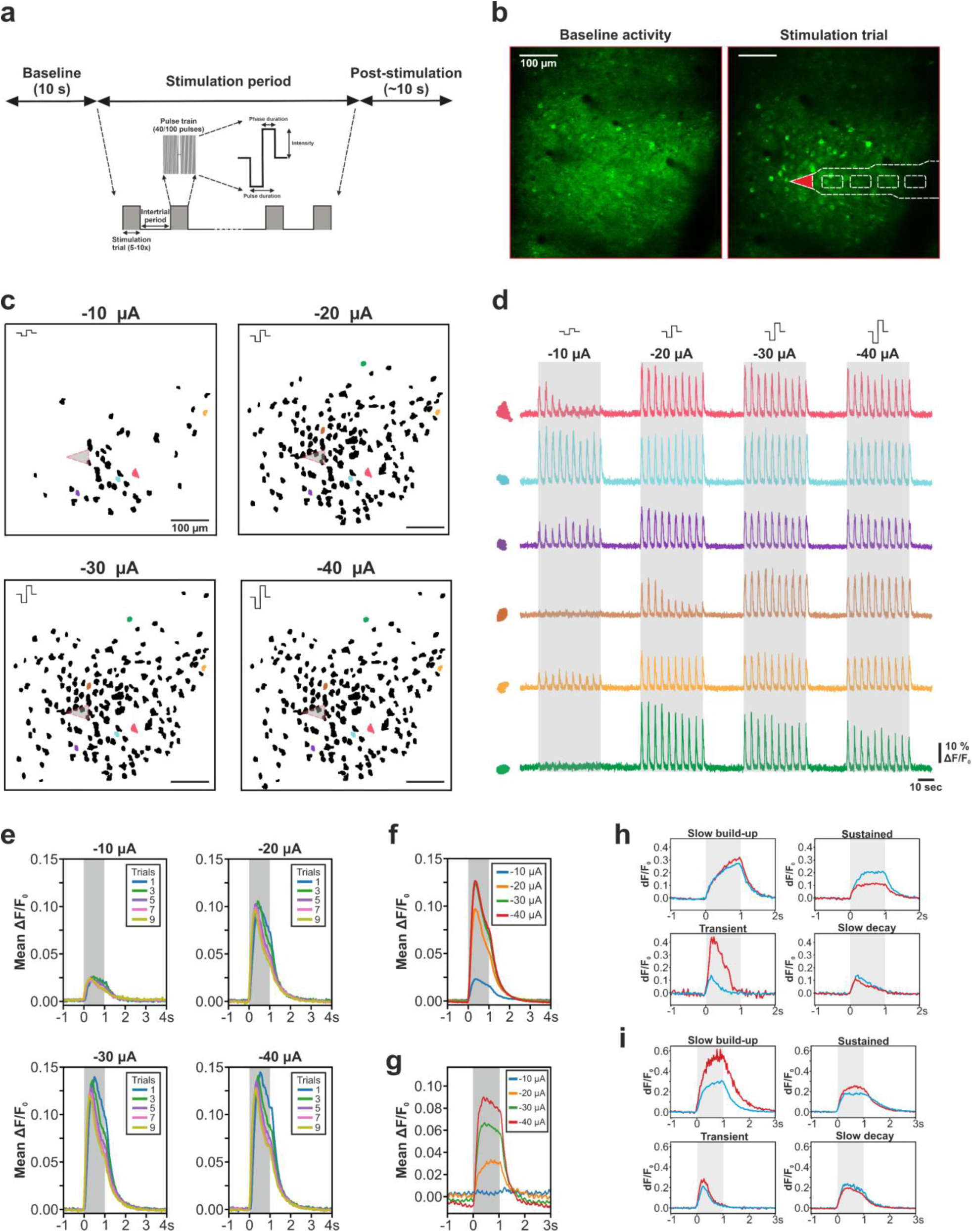
In vivo calcium imaging during monopolar electrical stimulation in anesthetized GCaMP6f mice. a) Schematic of the electrical stimulation protocol. b) Representative mean images (n = 30 frames) of the full field of view (FOV) during the baseline period (left) and during a stimulation trial (right). The schematic overlay on the right indicates the approximate position of one probe shank and the microelectrode used for stimulation (tip electrode, red). The probe was located below the imaging plane. c) Regions of interest (ROIs), corresponding to neurons activated by electrical stimulation, identified within the FOV at four different current intensities. Cathodic-first, biphasic pulses (n = 100) were delivered at 100 Hz with a pulse duration of 400 µs. The gray triangle marks the approximate location of the stimulation electrode. d) Calcium traces from six representative neurons recorded at different current intensities. The spatial locations of the neurons can be identified in panel c by the corresponding ROI shapes and colors. e) Mean calcium activity across all detected neurons at different current intensities, shown for odd-numbered stimulation trials. f) Mean calcium activity across all detected neurons at different current amplitudes, averaged across all stimulation trials. g) Same as panel f, shown for a different animal. h) Mean calcium traces from representative neurons exhibiting distinct response types to electrical stimulation. Two example neurons are shown for each response type. i) Same as panel h, shown for a different animal. In panels d-i, gray bars indicate the timing of stimulation trials.

Spontaneous neuronal activity in the superficial layers of sensory cortices is typically sparse in anesthetized rodents (Barth & Poulet, 2012; Petersen & Crochet, 2013; Sakata & Harris, 2009). Consistent with previous reports, only a small number of active neurons were detected during baseline periods, with activity increasing at lighter levels of anesthesia and becoming nearly absent during deep anesthesia (Figure 5b). In contrast, monopolar stimulation (1-s duration per stimulation trial) reliably evoked robust activation of neurons surrounding the stimulating electrode (Figure 5b), with the number of responsive cells increasing as a function of stimulation amplitude (Figure 5c). Higher current intensities also recruited neurons located at greater distances from the stimulation site, in agreement with earlier findings (Figure 5c; Lycke et al., 2023; Wu et al., 2023).

Neuronal calcium responses exhibited substantial heterogeneity across stimulation trials and current intensities (Figure 5d). For instance, at higher current amplitudes, the average calcium response (dF/F_0_) decreased over successive stimulation trials (Figure 5e), likely reflecting neuronal overstimulation or adaptation. Furthermore, as expected, the calcium response averaged across all neurons increased with current intensity (Figure 5f, g), reflecting the recruitment of additional neurons. The temporal profile of the Ca^2+^ response—particularly at higher currents—was characterized by a rapid onset immediately following the stimulation onset, peaking within a few hundred milliseconds, followed by a gradual decay (Figure 5f, g).

Based on calcium response dynamics, we could identify four distinct neuronal response types to electrical stimulation (Figure 5h, i). Three of these responses correspond to previously reported categories (Wu et al., 2023), and the observed response types also show substantial overlap with stimulation-evoked classes described in another study (Hughes et al., 2025). “Slow build-up” neurons exhibited a gradual and sustained increase in calcium signal throughout the stimulation period, while “sustained” neurons reached a plateau shortly after stimulation onset and maintained this level until stimulation offset. “Slow decay” neurons showed a gradual decline in calcium signal following an initial peaking, whereas “transient” neurons displayed a rapid decay, returning to baseline before the end of stimulation (Figure 5h, i). In the first three response types (“slow build-up, “sustained”, and “slow decay”), calcium levels typically returned to baseline within 0.5–1 s after stimulation cessation in GCaMP6f mice.

It is worth noting that in the Thy1-GCaMP6 mouse lines used in this study, the majority of labeled cortical neurons expressing the calcium indicator are excitatory pyramidal cells. To simultaneously record activity from inhibitory interneurons and excitatory neurons using calcium imaging, it is currently necessary to combine transgenic mouse lines with targeted viral injections (Dadarlat et al., 2024; Hughes et al., 2025).

To quantify the effects of monopolar stimulation, we analyzed both the number of neurons activated by stimulation and the mean calcium response across different current intensities, pulse durations, and stimulation frequencies in five animals (Figure 6; see Methods for details). As expected, both the number of activated neurons and the mean calcium response increased with increasing stimulation current (Figure 6a, d). On average, approximately 100 neurons were activated within a single imaging plane in layer 2/3 of V1 when a current of –40 µA was applied. The substantial inter-animal variability observed likely reflects differences in imaging quality, signal-to-noise ratio (SNR), and cortical vascularization across animals. For instance, although GCaMP6s mice exhibit a lower SNR compared to GCaMP6f mice, a higher proportion of neurons express the calcium indicator in GCaMP6s animals (Dana et al., 2014).

**Figure 6.**
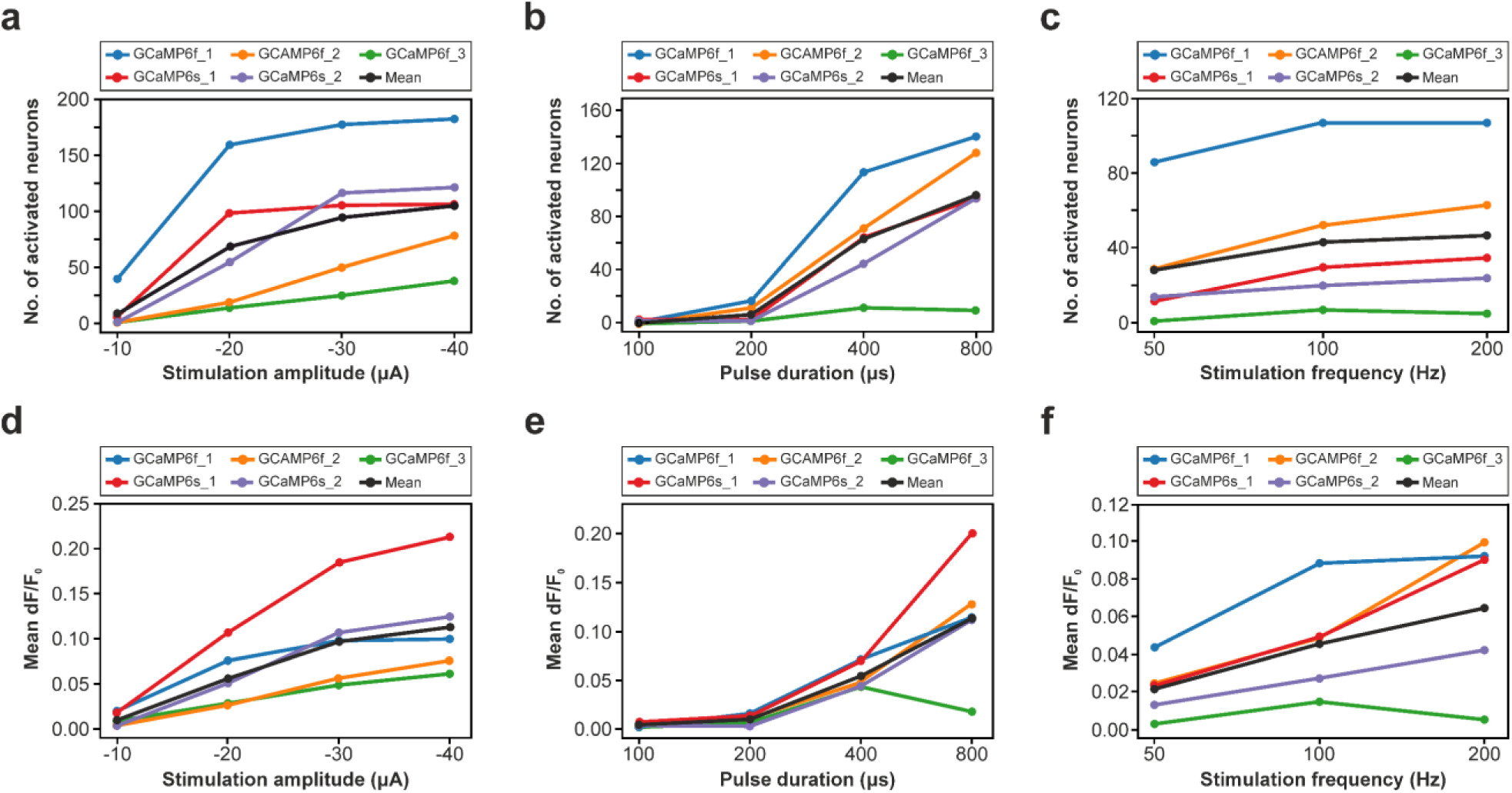
Quantitative comparison of the effects of monopolar stimulation parameters. a-c) Number of neurons activated by electrical stimulation as a function of current intensity (a), pulse duration (b) and frequency (c). d-f) Mean calcium response of stimulation-activated neurons as a function of current intensity (d), pulse duration (e) and frequency (f). Data from three GCaMP6f mice and two GCaMP6s mice are shown as colored lines. Population averages are indicated by solid black lines.

Furthermore, as anticipated, longer pulses activated more neurons and elicited larger calcium responses (Figure 6b, e). Very short pulses failed to evoke measurable activity, whereas longer pulses activated, on average, more than 80 neurons at moderate current amplitudes (−20 to −30 µA). In contrast, stimulation frequency had a comparatively modest effect on neuronal recruitment: the number of activated neurons was similar across the tested frequencies, with only a slight increase in calcium response amplitude at higher frequencies (Figure 6c, f). However, note that because the total number of pulses delivered per trial (n = 100 pulses) was kept constant across all frequency conditions, stimulation trials were shorter at higher frequencies.

We next examined the spatial organization of stimulation-evoked neuronal responses across different stimulation parameters. Specifically, we quantified the distance of activated neurons from the stimulating electrode and averaged these values in a GCaMP6f mouse (Figure 7a-c). As expected, the mean distance increased with higher stimulation intensities, longer pulse durations, and higher stimulation frequencies, indicating that progressively more distant neurons were recruited under stronger stimulation conditions (Figure 7a-c). This observation is consistent with the findings of previous reports (Dadarlat et al., 2024; Histed et al., 2009; Lycke et al., 2023; Wu et al., 2023). The rate of increase in distance closely mirrored the trends observed in Figure 6: pulse duration produced the steepest increase in recruitment distance, whereas stimulation frequency had a comparatively smaller effect.

**Figure 7.**
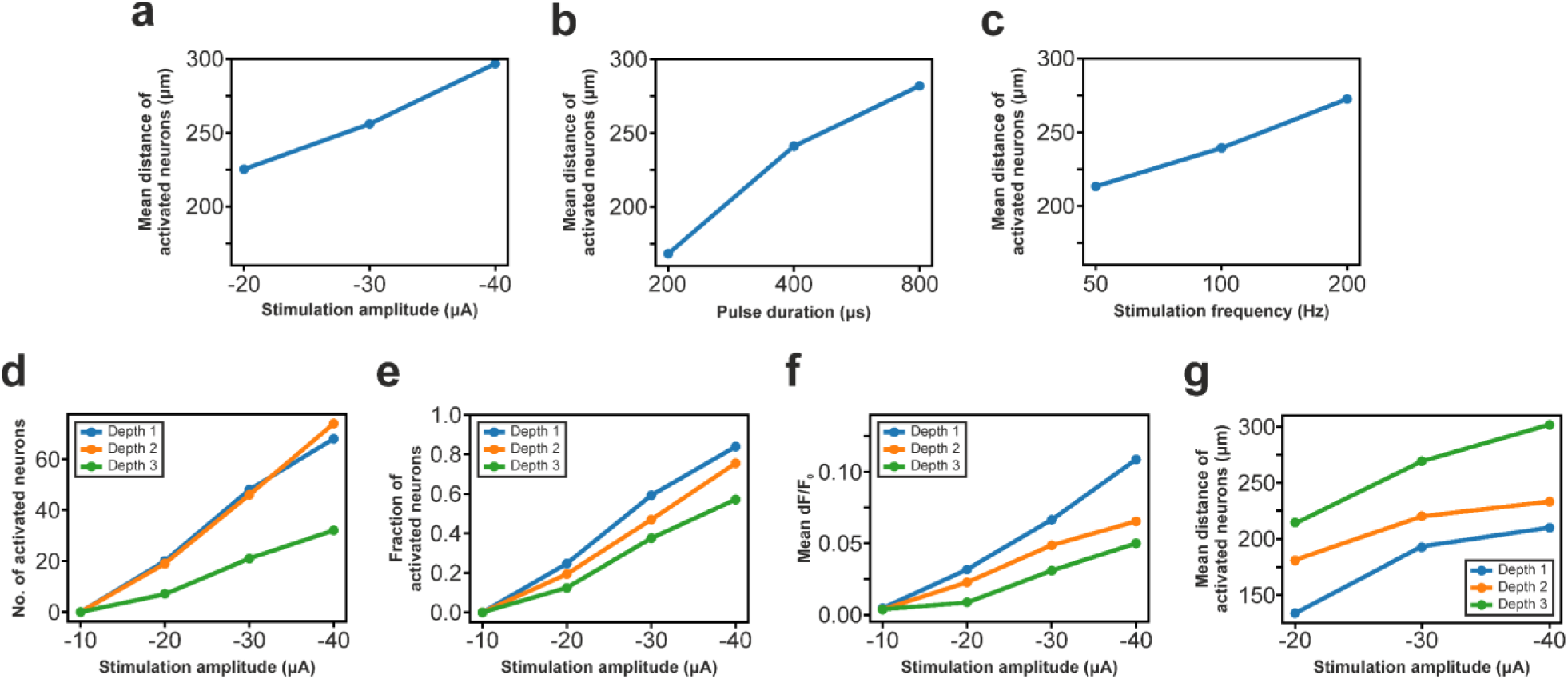
Distance- and depth-related effects of monopolar stimulation. a-c) Mean lateral distance of neurons activated by electrical stimulation from the stimulating electrode as a function of current intensity (a), pulse duration (b) and frequency (c). Data from a single mouse is shown. d) Number of neurons activated by electrical stimulation at different current intensities across three imaging depths separated by approximately 50 µm. The stimulating electrode was located closest to “Depth 1”, corresponding to the most superficial imaging plane. Data are shown for a single mouse. e) Same as panel d, but showing the fraction of activated neurons. f) Same as panel d, but showing the mean calcium response of activated neurons. g) Same as panel d, but showing the mean lateral distance of activated neurons from the stimulating electrode.

During resonant scanning, calcium signals are acquired from a single imaging plane. To assess the depth dependence of stimulation-evoked activity, we therefore performed consecutive recordings across multiple parallel imaging planes along the dorsoventral axis. In a GCaMP6f mouse, we performed calcium imaging during stimulation at varying current intensities in three imaging depths within layer 2/3, separated by 50 µm (Figure 7d-g). For each depth, we quantified the number and fraction of activated neurons (Figure 7d, e), the mean calcium response amplitude (Figure 7f), and the mean distance of activated neurons from the stimulating electrode (Figure 7g; only lateral distance from the electrode was considered for each depth). The stimulating electrode was located in the most superficial imaging plane (“Depth 1”).

Consistent with the lateral spread of activity observed earlier (e.g., see Figure 5c), both the number and proportion of activated neurons decreased with increasing distance from the stimulating site along the dorsoventral axis (Figure 7d,e). A similar depth-dependent decrease was observed in the mean calcium response (Figure 7f). These differences became more pronounced at higher current intensities. As expected, the mean distance of activated neurons increased in deeper imaging planes, reflecting the recruitment of fewer but more spatially dispersed neurons located farther from the stimulation site (Figure 7g). The observed properties of stimulation-evoked neuronal patterns are in agreement with previously reported findings (Dadarlat et al., 2024; Histed et al., 2009; Lycke et al., 2023).

### 2.5. *In vivo* calcium imaging in anesthetized mice during bipolar stimulation and current steering

While monopolar stimulation is one of the most widely used electrical stimulation strategies, it offers limited control over the spatial distribution and density of activated neurons (Moleirinho et al., 2021; Spencer et al., 2016; Zhu et al., 2012). More advanced stimulation strategies, such as bipolar stimulation or current steering, can enhance the spatial selectivity and resolution of neuronal activation; however, cortical responses to these patterns remain poorly understood (Berenstein et al., 2008; Bonham & Litvak, 2008; Dumm et al., 2014; Meikle et al., 2022; Zhu et al., 2012). The multi-shank probes and custom neurostimulator developed here enable simultaneous stimulation through multiple electrodes. To study stimulation-evoked cortical responses, the inter-shank spacing of the two-shank probe (650 µm) was chosen to allow the imaging FOV to be positioned between the shanks. This configuration enabled imaging within a cortical region largely free of implantation-related tissue damage and probe-induced fluorescence, as the probe shanks themselves often appear as bright fluorescent structures in the FOV.

In a GCaMP6f mouse, we applied bipolar stimulation between electrodes located on different shanks (Figure 8). In the first experiment, we selected electrode pairs vertically at different cortical depths, starting with the tip electrodes (Figure 8a). Although adjacent electrodes are separated by only 75 µm (center-to-center), the populations of neurons activated by different electrode pairs were largely distinct, showing minimal overlap across successive stimulation conditions (∼4%; Figure 8b). The number of activated neurons decreased with increasing distance from the imaged cortical area, and only a few neurons were activated by more than one electrode pair (Figure 8b).

**Figure 8.**
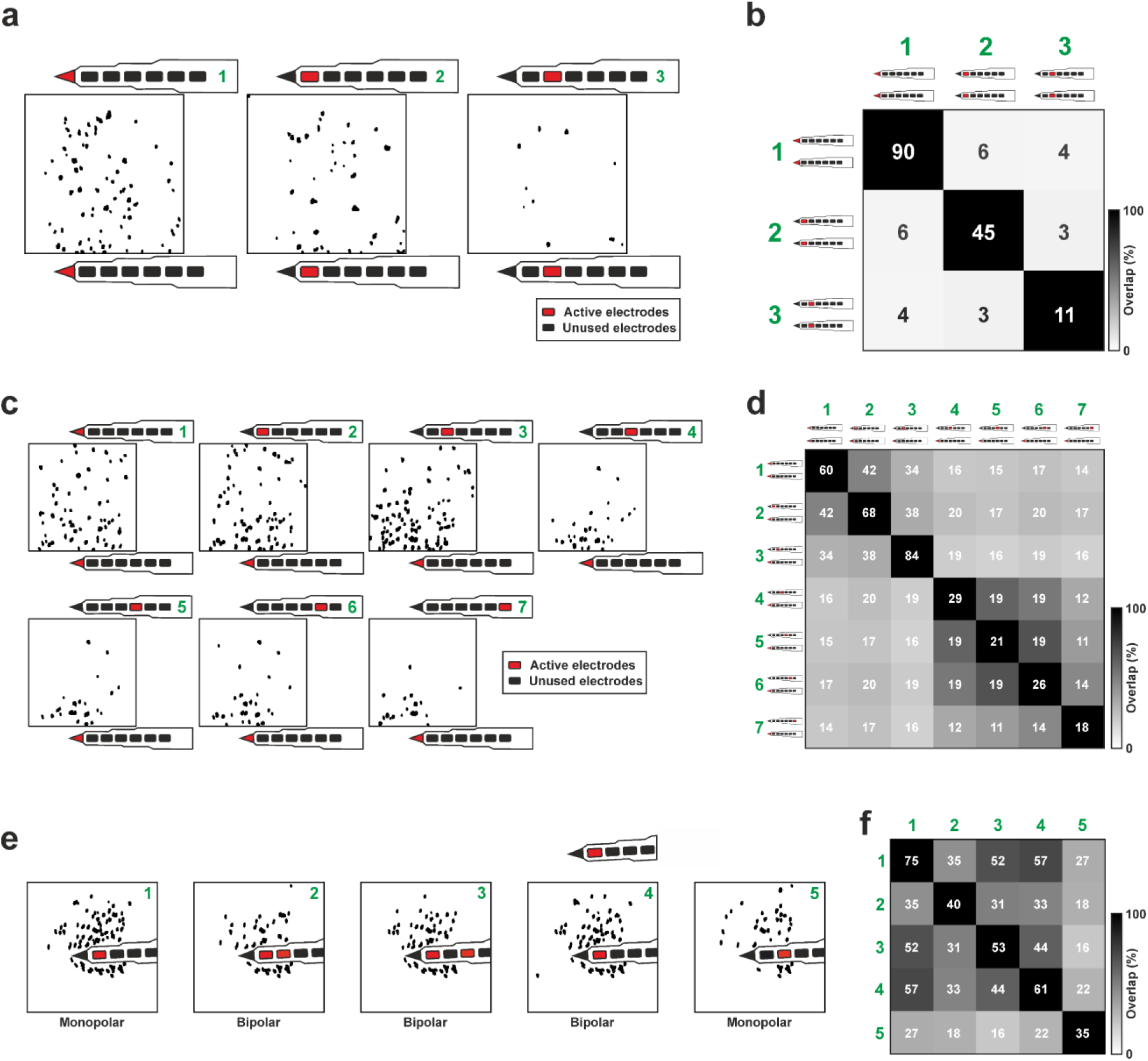
Calcium imaging during bipolar stimulation between probe shanks. a) Schematic of the experimental setup. The imaging field of view (FOV; black square) was positioned between the two probe shanks. The microelectrode pair used for stimulation is highlighted in red. Black patches within the FOV indicate the location of neurons activated by bipolar stimulation. Three distinct stimulation conditions were performed using different electrode pairs, with the imaging plane kept constant across all conditions. b) Grayscale map showing the overlap of activated neurons across the stimulation conditions depicted in panel a. Green numbers indicate condition identities. Overlaid numbers show the number of neurons activated in each condition (diagonal) and the number of neurons overlapping between conditions. c) Same as panel a, for a different animal. In this experiment, one electrode of the stimulating pair was held fixed while the other was varied across stimulation conditions. d) Overlap of activated neurons across the stimulation conditions shown in panel c. e) Distribution of neurons activated by monopolar and bipolar stimulation in another GCaMP6f mouse. Monopolar stimulation was delivered through two adjacent microelectrodes (conditions 1 and 5). Bipolar stimulation was performed using electrode pairs in which one electrode was the same as in condition 1, while the second electrode was either located on the same shank (conditions 2 and 3) or on the opposite shank (condition 4). f) Overlap of neurons activated across the stimulation conditions shown in panel e.

In the second experiment, one electrode of the pair (a tip electrode) was held fixed, while the second electrode on the opposite shank was systematically varied across stimulation conditions, starting from the shank tip (Figure 8c). In this configuration, the number of activated neurons again showed a decreasing trend as the second electrode was moved farther from the imaging plane, although the effect was less pronounced (Figure 8c, d). The overlap of activated neurons between stimulation conditions was substantially higher in this case (from ∼18% to ∼68%), likely because the fixed electrode consistently recruited nearby neurons across all stimulation conditions (Figure 8c, d). In contrast, the density of activated neurons in the upper region of the FOV - located closer to the variable second electrode - decreased as its distance from the imaging plane increased, with no neurons activated in this region during the final stimulation condition (Figure 8c).

In another experiment, we sequentially applied monopolar and bipolar stimulation at the same depth using identical stimulation parameters (−40 µA, 100 pulses, 400 µs pulse duration, 200 Hz frequency). One electrode of each bipolar pair was the same as that used for monopolar stimulation, and activated neurons were identified within the same FOV (Figure 8e, conditions 1–4).

When comparing stimulation conditions, monopolar stimulation activated the largest number of neurons within the FOV (condition 1). In contrast, bipolar stimulation using electrodes located on the same shank activated substantially fewer neurons (conditions 2 and 3). Under these conditions, activated neurons were more sparsely distributed and confined to a slightly smaller spatial area than during monopolar stimulation. Neuronal activation patterns were more similar to those observed during monopolar stimulation when the second electrode of the bipolar pair was positioned on the opposite shank (condition 4).

Interestingly, monopolar stimulation delivered through an electrode adjacent to the one used in condition 1 resulted in the lowest number of activated neurons (Figure 8e, condition 5), likely because this electrode was positioned slightly outside the imaging plane. The overlap of activated neurons across stimulation conditions was relatively high, ranging from ∼22% to ∼72% (Figure 8f). Together, these results support previous findings that monopolar and bipolar stimulation produce distinct spatial activation patterns, with bipolar stimulation using closely spaced electrodes generally eliciting more spatially confined neuronal responses (Schelles, Verhaege, et al., 2025; Uguz & Shepard, 2022; Zhu et al., 2012).

Next, we performed a current steering experiment in a GCaMP6f mouse using the two tip electrodes of the probe. The total stimulation current (50 µA) was divided between the two microelectrodes in 5 µA (10%) increments across 11 stimulation sessions, ranging from 0 µA to 50 µA on each electrode (Figure 9a). By progressively increasing the current on one electrode while decreasing it on the other, neuronal activation shifted systematically within the imaging field, with more neurons responding on the side of the FOV closer to the electrode carrying the higher current. In the two stimulation conditions where only one electrode was active (monopolar stimulation with 50 µA; Figure 9a, conditions 1 and 11), neuronal activation was largely confined to the side of the FOV near the active electrode, with only few neurons recruited on the opposite side. When equal currents were delivered to both electrodes (25 – 25 µA), activated neurons were distributed relatively evenly across the whole FOV (Figure 9a, condition 6). These findings are consistent with previous studies of current steering (Beauchamp et al., 2020; Dumm et al., 2014; Meikle et al., 2022).

**Figure 9.**
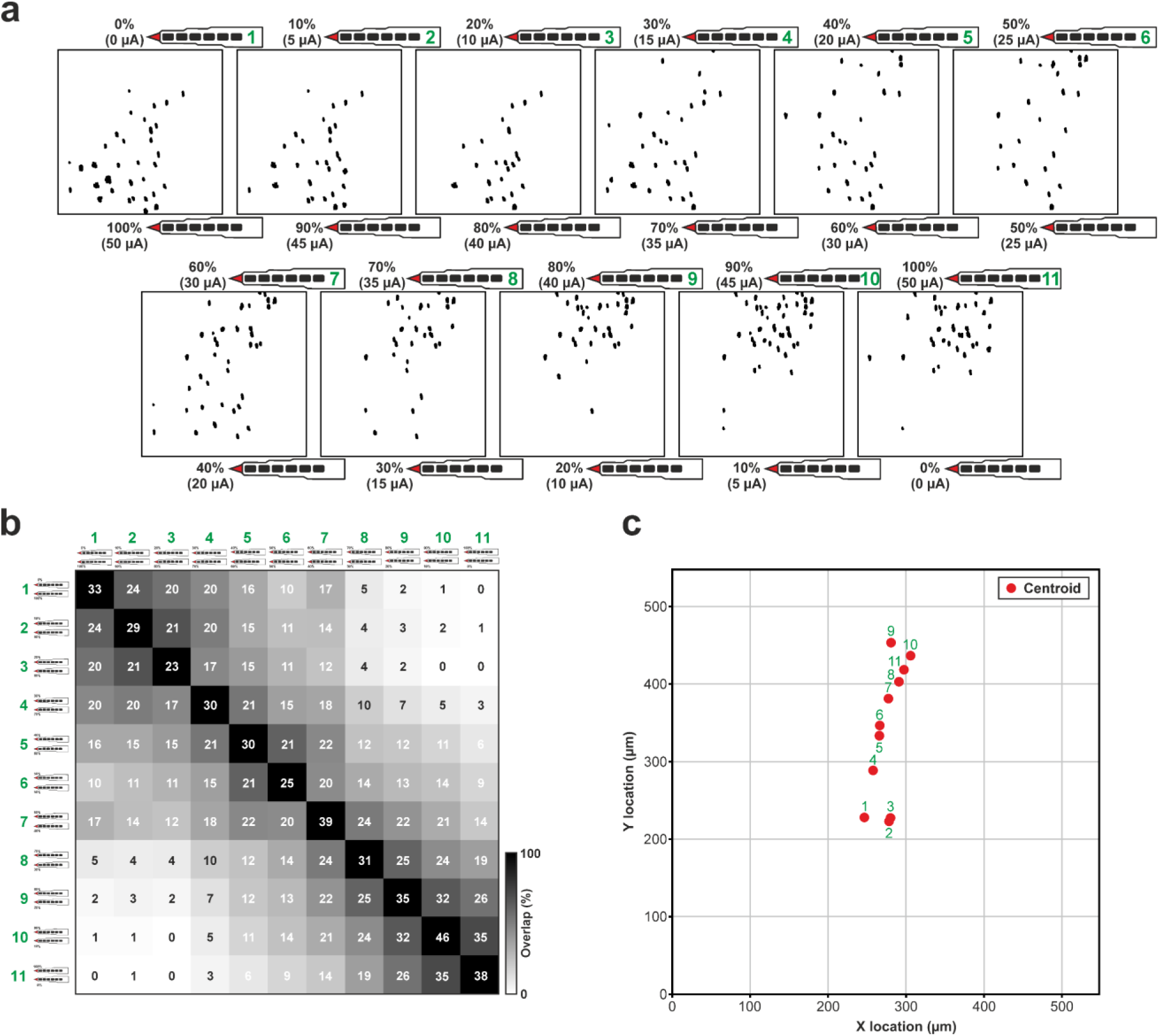
Calcium imaging during current steering between probe shanks. a) Schematic of the experimental setup. The imaging field of view (FOV; black square) was positioned between the two probe shanks. The microelectrode pair used for stimulation is highlighted in red. Current ratios and applied current intensities for each stimulation condition are indicated next to the FOV. Black patches show the locations of neurons activated by stimulation. b) Grayscale map showing the overlap of neurons activated across the stimulation conditions depicted in panel a. c) Centroid positions of all neurons activated under each stimulation condition, illustrating the systematic shift in neuronal activation across the FOV with changing current ratios. Green numbers indicate condition identities.

The total number of activated neurons remained relatively stable across stimulation conditions, ranging from 23 to 46. Overlap between activated neuron populations was highest when current ratios were similar between sessions (e.g., ∼63% between conditions 1 and 2) and lowest between the two monopolar conditions (0%), which recruited largely distinct neuron populations (Figure 9b). To further quantify these spatial effects, we computed the centroid of all activated neurons within the FOV for each stimulation condition (Figure 9c). The centroid shifted gradually and systematically from the middle of the FOV toward the upper section as the current ratio changed, moving from the side of the electrode delivering the higher current toward the opposite electrode (Figure 9c).

### 2.6. Simultaneous electrical stimulation and *in vivo* calcium imaging in an awake, head-fixed mouse

To investigate the effects of electrical stimulation under more physiologically relevant conditions, we implanted a flexible probe in a GCaMP6s mouse and performed simultaneous calcium imaging and electrical stimulation in the awake state (Figure 10). The probe was inserted into the visual cortex with its shank positioned beneath a glass coverslip (Figure 10a). After recovery, the animal was head-fixed in a custom-built setup, and monopolar stimulation was delivered through a rectangular electrode located directly above the tip electrode (Figure 10b).

**Figure 10.**
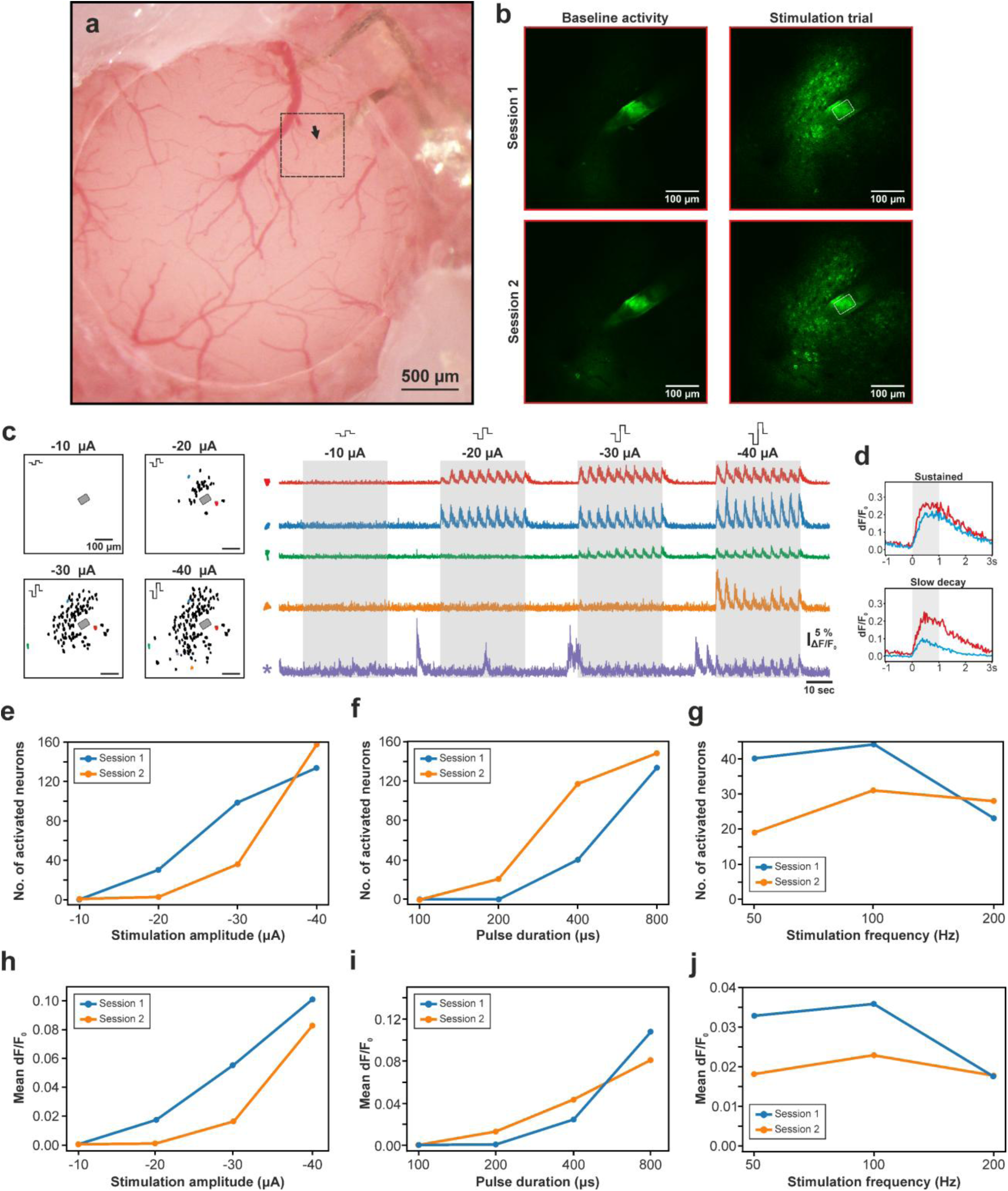
Calcium imaging in awake, head-fixed mice during concurrent electrical stimulation. a) Stereomicroscopic image of the craniotomy covered by a glass coverslip and the implanted probe (top right). The black rectangle indicates the field of view (FOV) shown in b. The arrow marks the approximate location of the microelectrode used for stimulation. b) Representative average images (n = 30 frames) of the full FOV during the baseline period (left) and during a stimulation trial (right) for two stimulation sessions. A section of the probe shank and the microelectrode used for stimulation are visible in the center of the FOV (outlined by a gray rectangle). c) Left: neurons activated by electrical stimulation within the FOV at four current intensities. Cathodic-first, biphasic pulses (n = 100) were delivered at 100 Hz with a pulse duration of 400 µs. The gray rectangle indicates the approximate location of the stimulation electrode. Right: calcium traces from five representative neurons across different current intensities. Spatial locations of neurons can be identified on the left by the shape and color of the ROI. The ROI marked by an asterisk indicates a neuron with strong spontaneous activity that was not classified as activated according to our criteria. d) Examples of neurons exhibiting different response types to electrical stimulation. Two representative neurons are shown for each type. Gray bars in panels c–d indicate the timing of stimulation trials. e-g) Number of neurons activated by electrical stimulation across different current intensities (e), pulse durations (f) and frequencies (g) during two imaging sessions performed on separate days. h-j) Mean calcium response of neurons activated by electrical stimulation across different current intensities (h), pulse durations (i) and frequencies (j) during the same two sessions.

During baseline periods, spontaneous activity near the stimulating electrode was relatively low but typically higher than under anesthesia (Figure 10b, c). As observed in anesthetized conditions, increasing the stimulation current recruited more neurons (Figure 10b, c). Interestingly, analysis of stimulation-evoked calcium responses of individual neurons revealed only two of the four response types identified in anesthetized mice - “sustained” and “slow-decay” - suggesting potential state-dependent differences in network dynamics (Figure 10d).

To assess the temporal stability of stimulation-evoked responses, we performed two stimulation sessions on separate days using identical stimulation parameters (the same as used for monopolar stimulation in anesthetized mice) and quantified both the number of activated neurons and the mean calcium response amplitude (Figure 10e-j). While the overall trends were similar between sessions, substantial variability was observed in absolute values, with the first session generally yielding stronger responses. This variability could arise from several factors, including differences in imaging quality (e.g., due to local inflammation or tissue swelling), slight probe displacement, variations in brain state, or minor differences in the imaging plane across sessions. Although image quality near the probe shanks declined a few days after implantation, likely due to local inflammation and thickening of the dura mater, robust stimulation-evoked calcium responses of individual neurons could still be observed in both V1 and higher-order visual areas even after several months, suggesting preserved functionality of the electrode array over chronic time scales (data not shown).

## 3. Conclusion

The findings of this study underscore the potential of flexible, polymer-based multi-shank probes as a robust platform for high-resolution neuromodulation and neuroscience research. By integrating MEMS-fabricated polyimide probes with a custom high-density neurostimulator, we have demonstrated a system capable of delivering complex electrical stimulation paradigms while simultaneously monitoring large-scale neuronal activity via two-photon calcium imaging. The mechanical flexibility, small cross-sectional area, and sharp-tipped design of the polyimide shanks markedly reduced tissue damage and brain dimpling, enabling high-quality imaging of superficial cortical layers that is often compromised when using traditional rigid electrodes.

Validation experiments in the mouse visual cortex demonstrated that monopolar stimulation produces reliable and repeatable activation of neuronal populations, consistent with previously reported findings and confirming the robust functionality of the platform. However, while monopolar stimulation effectively recruits neurons, it lacks the spatial precision required for sophisticated neural prosthetic applications. To address this limitation, we implemented and evaluated multielectrode stimulation strategies, including bipolar stimulation and current steering, which offer enhanced spatial selectivity and improved control over the electric field. We demonstrated that bipolar stimulation can activate neuronal populations distinct from those recruited by monopolar stimulation and can elicit more localized responses when the electrodes are placed in close proximity. Furthermore, our results with static current steering show that the centroid of neuronal activation can be finely tuned by adjusting current ratios between electrodes, allowing for precise control over the area of activation within the cortical tissue.

Although multielectrode stimulation protocols were implemented here using only two electrodes simultaneously, the neurostimulator system supports the concurrent selection of up to 16 electrodes, enabling the investigation of more complex stimulation paradigms, including tripolar and quadripolar configurations as well as dynamic current steering across multiple electrodes. This capability broadens the potential applicability of the stimulation platform across a wide range of neuromodulation applications. Furthermore, in this study we demonstrated two- and three-shank probes with 14 and 21 electrodes, respectively; however, the underlying microfabrication techniques readily allow these designs to be scaled, resulting in increased cortical coverage or higher electrode density through the addition of shanks or microelectrodes. Moreover, multiple electrode arrays can be interconnected to assess the effects of electrical stimulation in three dimensions. Finally, owing to their improved mechanical compliance with brain tissue, thin, flexible probes are expected to preserve tissue integrity and maintain stable stimulation performance over chronic timescales compared to rigid implants, making them a promising interface for implantable cortical sensory prostheses.

In conclusion, this integrated hardware and software platform, combined with two-photon calcium imaging, overcomes key limitations associated with electrical stimulation-induced electrophysiological artifacts and rigid neural implants. By enabling systematic characterization of local neural responses to advanced multielectrode stimulation paradigms, this approach may provide a powerful toolset for the development of next-generation sensory prostheses and clinical neuromodulation strategies aimed at eliciting more naturalistic and stable sensory percepts.

## 4. Experimental Section

### 4.1 Probe design

Custom electrode arrays having two or three shanks were fabricated from polyimide (Figure 1a,b). Each shank is 20 µm thick and 1000 µm long, with inter-shank pitches of 325 µm for the three-shank variant and 650 µm for the two-shank variant. Each shank contains seven iridium oxide electrodes: six rectangular microelectrodes (50 µm × 30 µm) arranged linearly, and a triangle-shaped electrode located at the shank tip. The microelectrodes have a center-to-center distance of 75 µm.

To facilitate handling of the device during surgery and implantation, a 75 µm-thick component made from SU-8, matching the dimensions of the probe base, was secured to the base with two-component epoxy glue (Figure 1b). The electrode array has a meandering cable to reduce tension forces during probe handling and implantation. A Samtec connector (CLP-111-02, Samtec) is soldered to the end of the flexible cable (Figure 1b) which can be connected to a Samtec-Omnetics adapter printed circuit board (PCB). The Omnetics connector (NPD-36-VV-GS, Omnetics Connector Corporation) on the adapter PCB allows direct connection to either the neurostimulator or the electrophysiological recording device.

### 4.2 Fabrication process

Figure 1c shows the steps of probe fabrication. First, a 10 µm thick polyimide base layer (PI2611, HD MicroSystems, USA) was spin coated and thermally cured at 200 °C, to increase the later adhesion with the top layer and thus the lifetime of the flexible implant (Ceyssens & Puers, 2015) (1). A platinum layer with a thickness of 200 nm (EVOCHEM Advanced Materials GmbH, Germany) was then sputter-deposited onto the polyimide and patterned using a lift-off process (2). To form the electrodes, iridium oxide was sputtered onto the platinum layer and again patterned with lift-off (Schelles, Goyvaerts, et al., 2025) (3). Sputtered iridium oxide is well suited for neural recording and stimulation due to its comparatively low impedance relative to conventional electrode materials such as gold or platinum. Subsequently, an additional 10 µm thick polyimide top layer was applied and baked at 350 °C, to fully cure the polyimide and ensure biocompatibility and a high longevity. The comb-like structures were defined by reactive ion etching, which opened up the electrode sites and bonding pads (4). In the final step, the 75 µm thick SU-8 reinforcement layer was fabricated by casting diluted SU-8 2050 (Micro Resist Technology, Germany) and deposited on top of the base, to facilitate handling and improve the mechanical stability of the probe (Merken et al., 2022) (5).

### 4.3 Neural stimulator

A custom neural stimulator was developed to accommodate the required stimulation patterns. Constant-current stimulation with a current resolution of 0.7 µA was delivered by four digital-to-analog converters (DAC8775, Texas Instruments, USA), with four analog output channels each. Six switch arrays (AD75019, Analog Devices, USA) were used to route these 16 independent stimulation channels to a total addressable amount of 96 electrodes. All electronic components were controlled by an Arduino Due microcontroller (Arduino, Italy) using a Serial Peripheral Interface (SPI) link. Before each stimulation trial, a custom Python script computed the necessary stimulation parameters, ensuring timely and accurate stimulation. Charge balancing was guaranteed by combining multiple techniques. First, each biphasic stimulation pulse was actively charge balanced using equal charges in the anodic and cathodic phase. During bipolar stimulation, in the absence of a ground electrode, the net current into the brain was always set to zero, meaning the sum of all cathodic currents had to equal the sum of all anodic currents, over all electrodes. Second, after each pulse, the electrodes involved in stimulation were briefly connected to the ground by means of the switch array, to get rid of any residual DC voltage across the electrodes.

### 4.4 Charge injection capacity measurements

To establish the level of safe stimulation, we performed *in vitro* charge injection capacity (CIC) measurements on the electrodes. This parameter reflects the maximum charge that can be safely delivered to the tissue during stimulation, while still preserving a safe voltage across the electrode-tissue interface, and thus without causing adverse effects. We determined the CIC by gradually increasing the current amplitude of each subsequent biphasic pulse in a pulse train, sent between a microelectrode and a large platinum/iridium return electrode, and measuring the corresponding voltage transients, using a third reference electrode. For each voltage transient, we calculated the voltage across the electrode-tissue interface as the capacitive part of the voltage transient, as this represented the actual charge buildup across the electrode-tissue interface. The current corresponding to a cathodic capacitive voltage drop of –0.6 V was considered the maximal safe current before hydrolysis of water would start to take place. The CIC was then calculated as this maximal safe current times the pulse width, divided by the area of the electrode (Boehler et al., 2020).

### 4.5 *In vitro* electrochemical impedance spectroscopy

Electrical impedance spectroscopy was performed on 77 microelectrodes across four probes following the procedure described in Horváth, Csikós, et al. (2024) (Figure 2a). Briefly, impedance magnitude and phase were measured using a nanoZ impedance testing device (White Matter LLC, USA) in 0.9 % NaCl solution, using an Ag/AgCl reference electrode, at thirteen biologically relevant frequencies (ranging from 1 Hz to 10 kHz). Microelectrodes showing impedance values above 1 MΩ at 1 kHz (indicating putative open circuits) were excluded from further analysis (n = 2 electrodes; 2.59 %). Before impedance measurement, the activation mode of nanoZ was applied to the microelectrodes using the default settings (activating current: from −1 µA to +1 µA, frequency: 1 Hz, duty cycle: 25 %, duration: 3 s, pause: 3 s).

### 4.6 Noise measurements

We used an Intan RHD-2000 electrophysiological recording system (Intan Technologies LLC, USA) to collect 3 minutes of *in vitro* recordings from probes (n = 4; two probes from each variant) submerged in 0.9 % NaCl solution. Data was acquired using a 32-channel headstage at a sampling rate of 20 kHz per channel with 16-bit resolution. A stainless-steel needle immersed in the NaCl solution served as the reference electrode.

Next, we computed the RMS noise level in the local field potential (1 – 500 Hz, 3rd-order Butterworth filter) and action potential (500 – 5000 Hz, 3rd-order Butterworth filter) frequency bands using the second minute of the *in vitro* recordings (Figure 2b). Values corresponding to high-impedant (putative nonfunctional) electrodes were excluded from analysis (n = 3 from a total of 70 microelectrodes). *In vivo* noise levels in the action potential band were estimated from cortical recordings obtained from an anesthetized mouse (n = 21 electrodes of a single probe; see Section “Processing and analysis of electrophysiology data” for details).

### 4.7 Animals

All experiments were conducted in accordance with the EC Council Directive of September 22, 2010 (2010/63/EU), and all procedures were reviewed and approved by the Animal Care Committee of the HUN-REN Research Centre for Natural Sciences and the National Food Chain Safety Office of Hungary (license number: PE/EA/01104-6/2022). Transgenic mice with genetically encoded intracellular calcium indicators were used for acute *in vivo* calcium imaging experiments, including GCaMP6s mice (n = 2; C57BL/6J-Tg(Thy1-GCaMP6s)GP4.3Dkim/J; Jackson Laboratories, Bar Harbor, ME, USA; RRID:IMSR JAX:024275) and GCaMP6f mice (n = 8; C57BL/6J-Tg(Thy1GCaMP6f)GP5.17Dkim/J; Jackson Laboratories, RRID:IMSR_JAX:025393) (Dana et al., 2014). One GCaMP6s mouse was used for awake, head-fixed experiments. C57BL/6J wild type mice (n = 2) were used for *in vivo* electrophysiology recordings. All experiments were performed on adult mice (>8 weeks of age). Animals were housed in 12-h light/dark cycles and had free access to food and water.

### 4.8 Animal surgery and probe implantation in acute experiments

Mice were anesthetized with a mixture of ketamine (100 mg/kg) and xylazine (10 mg/kg) administered intraperitoneally and updated regularly (1-2 injections per hour) to maintain a stable depth of anesthesia. Physiological body temperature of the animals was maintained by a homeothermic heating pad placed under them. The heating pad, equipped with internal temperature sensors, was regulated by a temperature controller unit (Supertech, Hungary).

After the animals reached the level of surgical anesthesia, they were positioned in a stereotaxic frame (David Kopf Instruments, USA), and the skin and connective tissue were removed to expose their skull. Next, a small craniotomy (2.5 mm x 2.5 mm) was prepared over the left primary visual cortex (V1) using a dental drill (anterior-posterior (AP): from –2.0 mm to –4.5 mm; medial-lateral (ML): from –1.0 mm to –3.5 mm; coordinates relative to bregma; Paxinos & Franklin, 2019). Before the surgery, mice received a subcutaneous injection of dexamethasone (0.05 ml; 2 mg/ml; Rapidexon, Dechra Veterinary Products, United Kingdom) to reduce brain edema in the craniotomy.

The flexible probe, held at the SU-8 support by a remote-controlled micro-tweezer toolset (MicroSupport Co., Ltd.; Japan), was mounted on a motorized stereotaxic micromanipulator (Robot Stereotaxic, Neurostar, Germany) and driven into V1 at an insertion rate of 2 μm/s and at a shallow insertion angle (∼55° from vertical; Figure 1d-f). To minimize brain dimpling during and after implantation, the dura mater was carefully pierced at the cortical entry point with a 34G bent needle.

The final target location and orientation of the probe shanks were determined based on the layout of surface vasculature. We typically targeted blood vessel-sparse regions to ensure good quality calcium imaging. The probe was advanced 700 µm into the tissue and then retracted by 100 µm, positioning the microelectrodes located at the bottom of the shanks within the superficial layers of V1 (layers 1-3), with the tip electrode located approximately 300 µm below the brain surface (Figure 1d). The probe was inserted with the microelectrodes facing ventrally.

For stabilization purposes, a thin layer of 1.5% agar or Dura-Gel (Cambridge NeuroTech, United Kingdom) was applied on the cortical surface, covering the implanted probe as well. A small amount of agar (or dental cement) was used to secure the meandering cable of the probe to the skull of the mouse. The micro-tweezer was then carefully detached from the probe and retracted. Up until the cortical surface was sealed with agar or Dura-Gel, room temperature physiological saline solution was regularly dripped into the craniotomy to prevent dehydration of the exposed brain tissue. Following probe implantation, the animal was transferred to the two-photon microscope for calcium imaging.

### 4.9 Animal surgery and probe implantation in the head-fixed mouse

All surgical procedures were performed under aseptic conditions. To implant the headplate and the flexible probe, anesthesia was induced using a mixture of intraperitoneally administered fentanyl (0.05 mg/kg), medetomidine (0.5 mg/kg) and midazolam (5 mg/kg). After the animal reached a surgical level of anesthesia, meloxicam (0.05 ml; 1 mg/kg; Melovem, Dopharma Research, Netherlands) and dexamethasone (0.03 ml; 2 mg/ml; Rapidexon) were administered subcutaneously, then the mouse was placed in a stereotaxic frame. Ophthalmic gel (Corneregel, Dr. Gerhard Mann Chem.-Pharm. Fabrik, Germany) was applied to keep the eyes moist throughout the surgical procedure. Body temperature was maintained at physiological levels using a homeothermic heating pad.

The scalp was disinfected with Betadine (Mundipharma, United Kingdom) and 70 % isopropyl alcohol, after which the skin and periosteum were removed to expose the skull. Next, a small amount of dental etching gel containing 38% phosphoric acid (Etch-Rite, PULPDENT Corporation, USA) was applied to the cleaned area to roughen the skull surface. This was followed by covering the skull with a thin layer of cyanoacrylate glue (Vetbond tissue adhesive; 3M, USA) to improve the adhesion of the dental cement applied later. After that, a rectangular craniotomy (∼4 mm x 3 mm) was prepared, centered over the left visual cortex. To keep the neocortex hydrated until the probe was implanted, sterile, room temperature physiological saline was regularly dripped into the cranial cavity.

Two circular glass coverslips with a diameter of 3 mm and 4 mm (Multi Channel Systems, Germany) were bonded together prior to surgery using optical adhesive (NOA 61, Norland Products, USA). The assembled coverslip was carefully placed in the anterior part of the craniotomy, with the smaller coverslip in direct contact with the cortical surface. Next, the coverslip was fixed to the skull using a small amount of light-curing dental cement (RelyX Universal Resin Cement, 3M).

The flexible probe was then implanted into the brain tissue using the same procedure as described for the acute experiments. The tips of the probe shanks, positioned in the posterior region of the craniotomy, were aligned near the edge of the coverslip and subsequently advanced beneath it. After the removal of the micro-tweezer, the part of the craniotomy not covered by the coverslip was sealed with a thin layer of Dura-Gel, followed by dental cement (Vertex Self Curing acrylic; Vertex Dental B.V., Netherlands). The connector of the probe was fixed to a small, flat stainless-steel support attached to the skull above the right hemisphere using the same type of dental cement. Finally, a custom stainless-steel headplate was secured to the anterior part of the skull with dental cement.

At the end of the procedure, anesthesia was reversed using a mixture of antagonists (atipamezole, 2.5 mg/kg; flumazenil, 0.5 mg/kg; naloxone, 1.2 mg/kg). Following surgery, the mouse was returned to the animal facility for a three-day recovery period, during which it received antibiotics (Augmentin, Glaxo Wellcome Production, France) and analgesics (Panadol Baby, Farmaclair, France). During calcium imaging sessions, the head of the mouse was secured using a custom head-fixation system.

### 4.10 Two-photon calcium imaging

Calcium imaging was performed using a Femto2D two-photon laser scanning microscope system (Femtonics, Hungary) equipped with a 20× water immersion objective lens (XLUMPLFLN; Olympus). The objective had a working distance of 2 mm, a numerical aperture of 1.0, and provided a field of view of 550 µm x 550 µm. Images were acquired at a resolution of 512 × 512 pixels using raster scanning at a frame rate of ∼31 Hz, corresponding to a spatial resolution of 1.08 µm × 1.08 µm per pixel.

A Coherent Chameleon Ultra titanium:sapphire femtosecond laser (Coherent, Saxonburg, USA) tuned to a wavelength of 920 nm was used for calcium imaging in GCaMP6f and GCaMP6s mice. After placing the mouse under the objective, the locations of the probe shanks and microelectrodes were first identified, after which intracortical microstimulation was started. Calcium imaging was typically performed at multiple imaging planes (i.e., cortical depths), depending on the positions of the microelectrodes used for stimulation. Approximately 0.5-2 hours of calcium imaging data were collected per animal. Acquired image files were stored on a network-attached storage device for subsequent offline analysis.

### 4.11 *In vivo* electrophysiological recordings

Acute *in vivo* neural recordings (n = 2 mice) were collected using an Intan RHD-2000 electrophysiological recording system (Figure 3). A 32-channel headstage equipped with an Omnetics connector was connected to the probe via the Samtec-Omnetics adapter PCB. We recorded spontaneous wideband cortical signals (0.1 – 7500 Hz) from V1 at a sampling rate of 20 kHz per channel with 16-bit resolution.

### 4.12 Electrical stimulation

Current-controlled electrical microstimulation was performed using a custom high-density neurostimulator device. The stimulator was controlled via Arduino and Python scripts, allowing flexible adjustment of stimulation parameters, including the number of pulses, current intensity and polarity, the duration and symmetry of biphasic pulses, stimulation frequency, the delay between consecutive stimulation trials, and the number of trial repetitions (for more details on the stimulator hardware and software, see Section 4.3, “Neural stimulator”). Electrical stimulation was triggered by a transistor-transistor logic (TTL) pulse generated by a PulsePal device (Sanders & Kepecs, 2014). Calcium imaging acquisition was synchronized to stimulation using the same TTL pulse. Each calcium imaging session started with a 10 s pre-stimulation (baseline) period, followed by the applied stimulation protocol (30-80 s), and ended with a 10 s post-stimulation period (Figure 5a). Electrode voltages were monitored simultaneously with calcium imaging using a PicoScope 2000 oscilloscope (Pico Technology, United Kingdom).

For validation experiments, monopolar stimulation was applied, in which one selected microelectrode on the probe served as the active electrode, while a stainless-steel needle with a large surface area placed on the cortical surface acted as the ground electrode. Stimulation consisted of charge-balanced, symmetric, biphasic pulse trains with the following default parameters: 100 pulses; total pulse duration (cathodic plus anodic phases) of 400 µs with no interphase interval; stimulation frequency of 100 Hz; inter-trial delay of 3000-5000 ms; and 5–10 trial repetitions. Current intensities of −10, −20, −30, and −40 µA were tested (Figures 5 and 10). In a subset of experiments, pulse duration (100, 200, 400, and 800 µs) and stimulation frequency (50, 100, and 200 Hz) were also systematically varied (Figures 6 and 10).

To assess the effects of the simultaneous use of multiple electrodes for stimulation on local cortical activity, bipolar stimulation (Figure 8) and static current steering (Figure 9) were employed. During bipolar stimulation, one active electrode delivered cathodic-leading pulses, while a second microelectrode simultaneously delivered anodic-leading pulses of equal current amplitude. Each stimulation trial consisted of pulse trains with current amplitudes of 10, 15, 20, 25, or 30 µA, delivered in randomized order at 8 s intervals. Each pulse train comprised 40 pulses applied at a frequency of 200 Hz. Stimulation trials were repeated six times, with 5 s inter-trial intervals. Results are shown only for 30 µA stimulation (Figure 8).

Another relevant electrical stimulation strategy is current steering, where we can focus the area of neuronal activation by varying the ratio of the current amplitude between two (or more) electrodes. Current steering was implemented using a static paradigm, in which the ratio of current amplitudes between two electrodes remained constant throughout each stimulation condition. In these protocols, pulse polarity was identical across the simultaneously stimulated electrode pair. Static current steering was applied between the two tip electrodes, with the total injected current summing to 50 µA (Figure 9). A stainless-steel needle placed on the cortical surface served as the ground electrode. Current amplitudes were varied in 5 µA increments between 0 and 50 µA, resulting in 11 distinct stimulation conditions.

For bipolar stimulation and current steering experiments, overlap of activated neurons between stimulation conditions (shown as grayscale maps in Figures 8 and 9) was quantified as the number of neurons activated in both conditions divided by the total number of unique neurons activated across the two conditions. The overlap is expressed as a percentage.

### 4.13 Histology

Nissl-staining was performed to identify the tracks of the polymer probes in brain tissue and to verify the recording locations, similarly as described in Horváth, Ulbert et al. (2024) (Figure 1f). In brief, following the acute experiments, animals were deeply anesthetized with a high dose of ketamine/xylazine and were perfused through the heart, first with 50 ml of physiological saline, followed by 150 ml of 4% paraformaldehyde in 0.1 M phosphate buffer (pH = 7.4) to fix the brain tissue. Following perfusion, the brain was removed from the skull, post-fixed in the same fixative solution, and stored at 4°C until further histological processing.

Sagittal brain sections (60 µm thick) were cut using a vibratome (Leica VT1200, Leica Microsystems, Germany). Sections were then washed in 0.1 M phosphate buffer and transferred to a gelatin-filled Petri dish. From there, the brain slices were carefully mounted onto microscope slides and air dried. After that, cresyl violet (Nissl) staining was applied on the sections, followed by dehydration in xylene. Slides were then coverslipped using DePex (SERVA Electrophoresis, Germany). Finally, Nissl-stained sections containing probe tracks were photographed under an optical microscope (Leica DM2500, Leica Microsystems; <u>RRID:SCR_020224</u>) equipped with a digital camera (DP73, Olympus, Japan).

### 4.14 Processing and analysis of calcium imaging data

Suite2p (v0.14.3; <u>RRID:SCR_016434</u>) was used to identify regions of interests (ROIs, i.e., putative neurons) in the calcium imaging data (Pachitariu et al., 2017). In most cases, default parameters were used for motion correction and ROI detection. However, in some measurements, the spatial scale and threshold scaling parameters were adjusted to improve detection accuracy. The decay time (tau) was set to 0.5 s for GCaMP6f-expressing mice and to 1.25 s for GCaMP6s-expressing mice. The outputs of suite2p were then visually inspected and manually curated to refine ROI selection. Specifically, artifactual ROIs (e.g., those detected next to the probe shank) were removed, or additional ROIs corresponding to neurons missed by the automated algorithm were manually added.

Next, for each ROI, we calculated the baseline fluorescence intensity (F_0_) as the average calcium signal during the pre-stimulation period (with a typical duration of ∼10 s). Then we computed the baseline-corrected calcium trace (dF/F_0_) for each ROI. To identify neurons activated by electrical stimulation, we compared the mean calcium activity during the stimulation period with baseline activity for each neuron. A response threshold was defined as the mean baseline signal plus a multiple of its standard deviation, with the multiplier adjusted according to the signal-to-noise ratio of the calcium imaging data. Neurons exceeding this threshold during any stimulation trial were classified as activated (e.g., see Figure 5c).

For each stimulation condition, we calculated the number of activated neurons and the mean calcium response (dF/F_0_) across all identified neurons. Furthermore, to characterize the temporal dynamics of stimulation-evoked responses, dF/F₀ traces spanning several seconds were computed for each stimulation condition and averaged across trials or neurons, as appropriate. In a subset of experiments, the distance between the center of the active microelectrode and the center of each ROI was also determined to assess changes in the mean distance of activated neurons across stimulation conditions.

For current steering experiments, we calculated the centroid of the activated neuron population to visualize shifts in activity within the imaging plane between stimulation conditions using different current ratios (Figure 9c). Finally, ROIs corresponding to neurons activated by the applied stimulation protocols were plotted separately for each stimulation condition, and the overlap of activated neuron populations across stimulation conditions was assessed (Figures 8 and 9). Overlapping ROIs were identified using CellReg (v1.5.9; Sheintuch et al., 2017).

### 4.15 Processing and analysis of electrophysiological data

To isolate the activity of single neurons, we applied spike sorting as described in Horváth, Csikós et al. (2024). Kilosort2 was used to extract single-unit activity (Pachitariu et al., 2024; Pachitariu et al., 2016), then Phy2 (https://github.com/kwikteam/phy) was applied for manual curation of resulting single unit clusters. Clusters exhibiting atypical spike waveform shapes or clear noise contamination were excluded from further analysis. Additional exclusion criteria included a low firing rate (< 0.05 Hz, or fewer than 100 spikes) and a highly contaminated refractory period, as indicated by the autocorrelogram.

Next, for each isolated single unit, we calculated the average multichannel spike waveform from the wideband recording. We then computed the trough-to-peak amplitude of the mean spike waveforms, defined as the difference between the most negative trough and the largest positive peak of the spike waveform, measured on the channel exhibiting the maximal spike amplitude.

To estimate the noise level in the frequency band between 500 and 5000 Hz from *in vivo* recordings (for n = 21 microelectrodes of a single probe), we computed the RMS value for each electrode within a 50 ms windows centered on the inactive phases (down-states) of cortical slow waves induced by ketamine/xylazine anesthesia. During these inactive phases, cortical neurons cease firing, enabling reliable estimation of electrical noise levels *in vivo* (Fiath et al., 2016). Only down-states lasting at least 200 ms were included in the analysis, and RMS values were averaged across down-states. Down-states were detected using a multi-unit activity–based method, following previously described approaches (Fiath et al., 2016; Horváth, Ulbert, et al., 2024).

### 4.16 Statistical analysis

Box-whisker plots (Figures 2b and 3d) illustrate the distribution of the data as follows: the central line denotes the median, the box spans the 25th to 75th percentiles, and the whiskers indicate the minimum and maximum values. The mean is shown as a black dot, while individual data points are represented by smaller colored dots.

## Acknowledgements

This project has received funding from the European Union’s Horizon Europe research and innovation programme under the European Innovation Council (EIC) Pathfinder grant agreement No. 101071015 (HYPERSTIM). Project no. 150574 has been implemented with the support provided by the Ministry of Culture and Innovation of Hungary from the National Research, Development, and Innovation Fund, financed under the STARTING_24 funding scheme. R.F. was supported by the Bolyai János Scholarship of the Hungarian Academy of Sciences. E. N. was supported by the EKÖP-24-1 University Research Scholarship Program (EKÖP-24-1-PPKE-44) of Ministry for Culture and Innovation of Hungary from the National Fund for Research, Development and Innovation. The research leading to these results has received funding from the Hungarian Brain Research Program Grant (NAP2022-I-2/2022).

## Author contributions

**Eszter Nguyen** (Data curation, Formal analysis, Methodology, Software, Validation, Visualization, Writing— original draft, Writing—review & editing), **Balázs Barkóczi** (Investigation, Data Curation, Writing—review & editing), **Csaba Horváth** (Investigation, Data Curation), **Linda Judák** (Resources, Writing—review & editing), **Balázs Rózsa** (Resources), **Frederik Ceyssens** (Conceptualization, Methodology, Resources, Visualization, Writing—review & editing), **István Ulbert (**Funding acquisition, Supervision, Writing—review & editing), **Lucia Wittner** (Conceptualization, Supervision, Writing—review & editing), **Maarten Schelles** (Conceptualization, Methodology, Resources, Software, Visualization, Writing - Original Draft, Writing—review & editing), **Richárd Fiáth** (Conceptualization, Data curation, Investigation, Methodology, Supervision, Funding acquisition, Project administration, Writing— original draft, Writing—review & editing)

## Conflict of Interest

F. C. and M. S. are affiliated with ReVision Implant (Belgium), which develops technology for cortical visual prostheses.

## Data Availability Statement

The data that support the findings of this study are available from the corresponding author upon reasonable request.

